# Novel Glomeromycotina-Moss Associations Identified in California Dryland Biocrusts

**DOI:** 10.1101/2024.12.27.630416

**Authors:** Kian H. Kelly, Claudia Coleine, Chris Coshland, Jason E. Stajich

**Author notes:** Address correspondence to: Kian H. Kelly,; Jason E Stajich.

## Abstract

Drylands, which comprise ~45% of Earth’s land area, host biological soil crusts (biocrusts)—symbiotic communities of cyanobacteria, fungi, algae, lichen, and bryophytes that stabilize soil and support key ecosystem functions. Moss dominated biocrusts in particular are interesting due to their potential to illuminate ancient bryophyte-fungal interactions, but their associated microbial communities remain poorly characterized. To test the hypothesis that mosses in biocrusts harbor specific fungal associates, we conducted amplicon metabarcoding and microscopic surveys employing fungal staining across an aridity gradient. We identified novel associations between mosses and arbuscular mycorrhizal fungi (AMF), with phylogenetic analyses revealing distinct fungal communities in moss biocrusts compared to adjacent bare soil. Intracellular branching by fungi resembling Glomeromycotina was observed within healthy *Trichostomopsis australaceae* cells, suggesting interactions beyond saprotrophy. Moreover, shifts in AMF community composition along an aridity gradient highlight the potential vulnerability of biocrust mycobiomes to climate change. These findings provide critical insights into ancient bryophyte-fungal symbioses, potentially analogous to those enabling early land plant colonization during the Ordovician (~470 Ma). They also underscore the need to understand and protect biocrust microbial communities as aridity intensifies under climate change.

## Introduction

Drylands are biomes at the cusp of life and death, where extreme adapted microorganisms persist and, unbeknownst to many, provide critical services to the planet. Drylands encompass approximately 45% of Earth’s terrestrial surface and are expected to increase in aridity and size as climate change progresses (Prăvălie et. al 2016, Coleine et. al 2024). This effect is particularly pronounced in the Western United States, where water shortage will exceed the most severe megadrought periods of the last millennium (Cook et al. 2015). Biological soil crusts (hereafter, biocrusts) cover up to 70% of drylands (Barger et. al 2006, Elbert et. al 2012). Biocrusts are symbiotic communities of cyanobacteria, microfungi, algae, lichen, and bryophytes which aggregate and form a “crust” that occupies the top few millimeters of soil.

Biocrusts are crucial remediators of climate change. The global net carbon uptake of biocrusts from the atmosphere (photosynthesis minus respiration) is roughly ~3.9 gigatons per year. This represents approximately half the global annual carbon release from fossil-fuel combustion (~7.0 gigatons per year) (Elbert et. al 2012). Biocrusts perform other critical ecosystem services including soil stabilization, nutrient cycling, soil fertilization, and vascular plant growth promotion. They have been identified as an indicator of general ecosystem health in drylands (Weber et al. 2016, Bowker et al. 2018). The ecosystem services provided by biocrusts are driven, in part, by the physical and biochemical activity of the diverse microbes that comprise them (Barger et. al 2006, Elbert et. al 2012).

Previous studies have quantified the contributions of fungal and bacterial biocrust constituents to dryland nitrogen and carbon cycling (Zhao et al. 2020, Tian et al. 2023). They have also determined microbial physical contributions to biocrust stability, which helps prevent erosion and dust storms (Weber et al. 2016, Chamizo et al. 2017). Although biocrusts are key climate change remediators, climate change also puts biocrusts, and biocrust microbes, at risk. Biocrusts are predicted to experience up to 39% total land area loss by 2070 (Rodriguez-Caballero et al., 2018). This effect is likely to create a positive feedback loop which results in significant dryland degradation and climate change acceleration. Despite its clear importance, few studies have characterized the impacts of climate change on biocrust microbial communities.

Although several studies have characterized bacterial communities within biocrusts, relatively little work has focused on the biocrust mycobiome. There are several biocrust types, each with variable fungal and bacterial constituents (Pombubpa et al. 2020). These include lichen crust, cyanobacterial crust, and moss crust among others. Moss biocrusts are particularly susceptible to climate change (Ferrenberg et al. 2015). This may be, in part, due to shifts in the moss mycobiome (including symbiotic partners) caused by altered ecosystem water availability and temperature. The biocrust mycobiome may experience a shifted fungal repertoire at higher aridity, thereby altering the crust’s successional stage and ecological function (Ferrenberg et al. 2015, Pombubpa et al. 2020).

In addition to their critical ecosystem functions, biocrusts provide a unique opportunity to study interactions between early divergent plants (Bryophyta, e.g., mosses) and early divergent fungi. These interactions mirror those that facilitated the colonization of land by plants from aquatic origins (Hernández-Hernández et al. 2017, Bi et al. 2021, Field et al. 2015). The colonization of terrestrial environments by plants transformed Earth’s surface into a habitat capable of supporting more complex life forms and ecosystems (Schreiber et al. 2022).

Ancestral arbuscular mycorrhizal fungi (AMF; Glomeromycotina) are widely regarded as the earliest symbiotic partners of land plants (Strullu-Derrien et al. 2019, Field et al. 2015). Evidence supporting this includes the presence of genes required for AMF symbiosis in all land plants (Radhakrishnan et al. 2020, Wang et al. 2010) and fossilized arbuscules in healthy cells of early land plants (Strullu-Derrien et al. 2007, Strullu-Derrien et al. 2019). Interestingly, mosses (Bryophyta) are the only major lineage of land plants thought to lack fungal symbioses (Field et al. 2015). This contrasts with other bryophytes, such as liverworts and hornworts, which form AMF symbioses. Whether fungi in moss biocrusts establish symbiotic relationships with mosses remains unclear, but confirming such interactions could significantly enhance our understanding of both biocrust fungal ecology and ancient plant-fungal associations.

To date, direct evidence of nutrient exchange between mosses and fungi remains unpublished, and fungal hyphae have not been conclusively observed in symbiosis with healthy moss cells. This has perpetuated the paradigm that mosses, with the exception of the genus *Takakia*, are the only major lineage of land plants that do not form fungal symbioses (Radhakrishnan et al. 2020, Wang et. al 2010, Satjarak et al. 2022). However, this assumption may be premature, as only a limited number of moss species have been surveyed. The discovery of Glomeromycotina symbioses in mosses would further support the hypothesis that AMF were among the earliest symbionts of land plants, potentially overshadowing other candidates, such as Mucoromycotina.

Given the extensive evolutionary history of mosses and fungi, spanning hundreds of millions of years, it is unlikely that no symbiosis has evolved. Indeed, some mosses rely on co-opted genes from Mucoromycota fungi for key developmental processes such as stem cell and gametophore formation (Wang et al. 2020, Wang et al. 2021). Despite this, interactions between early divergent fungi and mosses remain understudied. Few microscopic surveys have visualized fungal structures in mosses from natural environments (Zhang and Guo 2007, Valdés et al. 2023), and even fewer have investigated physiological interdependence (Mathieu et al., 2024; Chen et al., 2022; Hanke and Rensing, 2010). This paucity of data is surprising, considering the ecological importance of mosses, their species richness (>10,000), and their potential to illuminate the evolution of plant terrestrialization (Field et al., 2015).

Here, we survey the associations between moss biocrusts and early divergent fungi across an aridity gradient in Southern California. First, we hypothesized that mosses are hosts for endophytic early divergent fungi. Secondly, that local climate would impact the recovery of early divergent fungi in our survey. To this end, we employed a dual metabarcode sequencing and microscopic survey, using staining to observe fungal structures in mosses collected from drylands.

The survey reveals novel associations between mosses and AMF. It also reveals a broadly distributed arbuscular mycorrhizal moss mycobiome across the Mojave Desert, Colorado Desert, and California coast in cosmopolitan moss *Trichostomopsis australaceae*. This moss mycobiome was distinct from the adjacent bare soil. Staining of moss tissue revealed intracellular branching of cells that are morphologically similar to Glomeromycotina within healthy *T. australaceae* cells, suggesting fungal-moss interactions beyond saprotrophy. Shifts in moss-associated Glomeromycotina composition along an aridity gradient within the same host highlight the potential impact of aridity on the biocrust mycobiome. This survey also offers key insights into bryophyte-fungal associations which may resemble those that enabled the colonization of land by plants.

## Materials and Methods

### Collection of moss biocrusts

Moss biocrusts were collected from two sites in the Mojave Desert, two sites in the Colorado Desert, and two coastal sites along a Northeastern transect. Cima Volcanic Field (CIMA, GPS: 35.1997917 N, 115.8699722 W) is dominated by volcanic rock and situated in the central Mojave Desert. Granite Mountains (GMT, GPS: 34.7804861 N, 115.6297361 W) is dominated by granitic soil and located in the Southern Mojave Desert. The Mojave Desert is known for its extreme hot-cold temperature fluctuations and slightly higher elevation than the Colorado Desert (Bai et al, 2014). All Mojave Desert sites were hyper arid (table 1). Oasis De Los Osos Reserve (ODLO, 33.89182 N, 116.68908 W) is located in the Northern Colorado Desert, while Anza Borrego Research Center (AB, 33.2407889 N, 116.3895333 W) is located in the Western Colorado Desert. The Colorado Desert experiences less extreme temperature fluctuations and higher water availability (Weiss and Overpeck 2005). All Colorado Desert sites were classified as arid (table 1). Mixed woody and succulent scrub dominated all sites, with Joshua Trees and Yucca plants visible at the Granite Mountains. Torrey Pines State Preserve is located on the coast of the Pacific Ocean and known for its maritime climate, large sandstone bluffs, and rich biodiversity. It is classified as semi-arid (table 1). Two sites were selected: one a sandstone bluff without proximal vegetation (TP, 32.9456861 N, 117.2494806 W), the other on sandy soil with woody shrubs within three feet (TP2, 32.9421506 N, 117.2482757 W). Smooth moss biocrusts with the same morphotype (dark, short moss) were collected opportunistically using sterile technique from each site in triplicate, as was the nearest bare soil (within three feet). All collected mosses were healthy and possessed green tissue (Figure 2). A metal spatula was thoroughly sterilized with 70% ethanol between the collection of each sample. The spatula was used to pry up biological soil crusts or scoop dirt into 50 mL falcon tubes or sterile bags. Samples were stored in a cooler of dry ice and transported to University of California Riverside and stored at −20°C.

**Figure 1:**
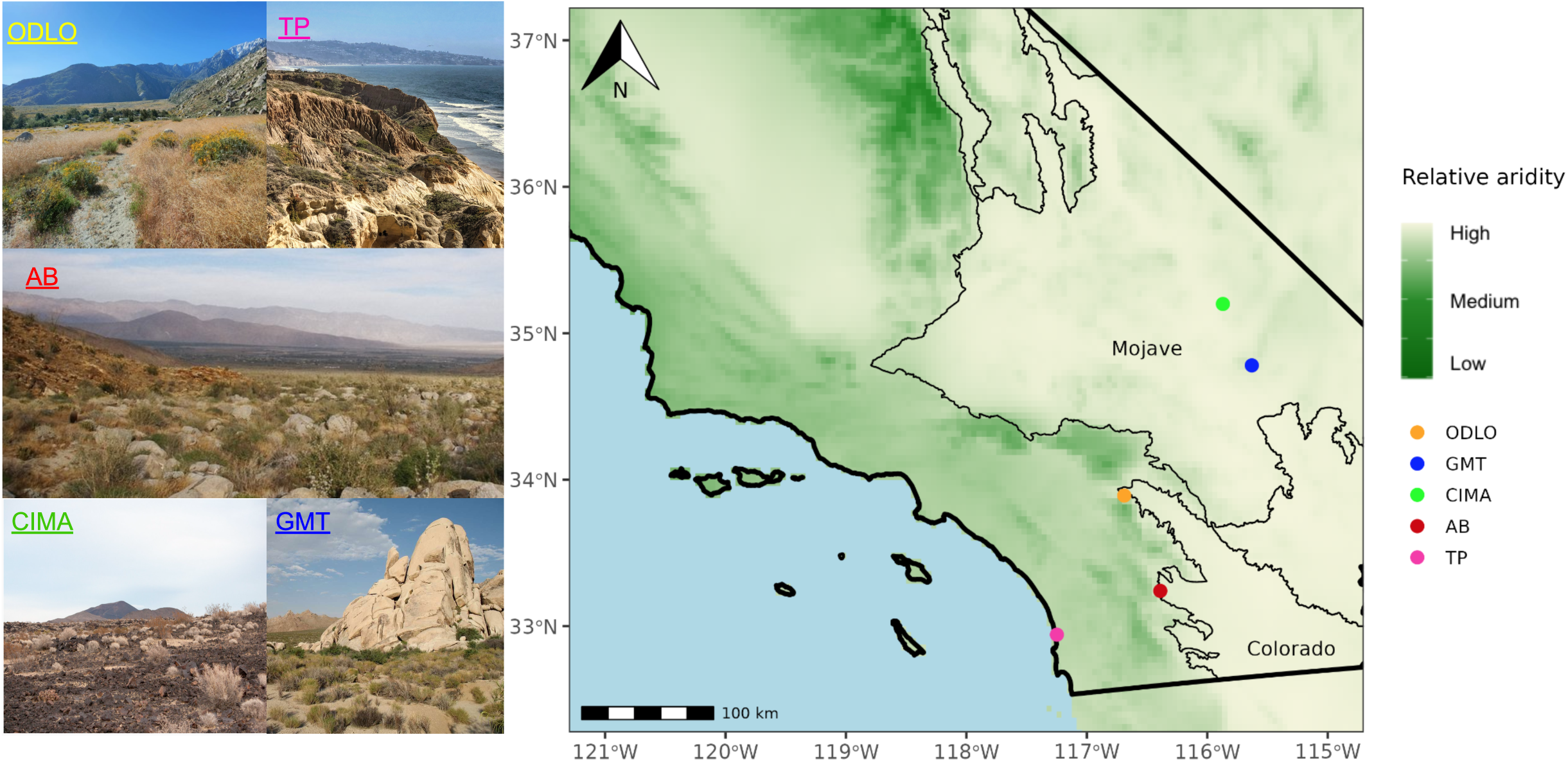
Sampling sites displayed on a map of relative aridity. Biological soil crusts dominated by healthy moss of the same morphotype were collected in triplicate across an aridity gradient along a Northeast transect. Aridity was calculated using the TerraClimate dataset and log normalized for comparison of sites (Abatzoglou et al. 2018). TP = Torrey Pines, ODLO = Oasis De Los Osos Reserve, AB = Anza Borrego Research Station, CIMA = CIMA Volcanic Field, GMT = Granite Mountains Research Center. Borders of the Mojave and Colorado Desert obtained from the US Geological Survey are displayed.

**Figure 2:**
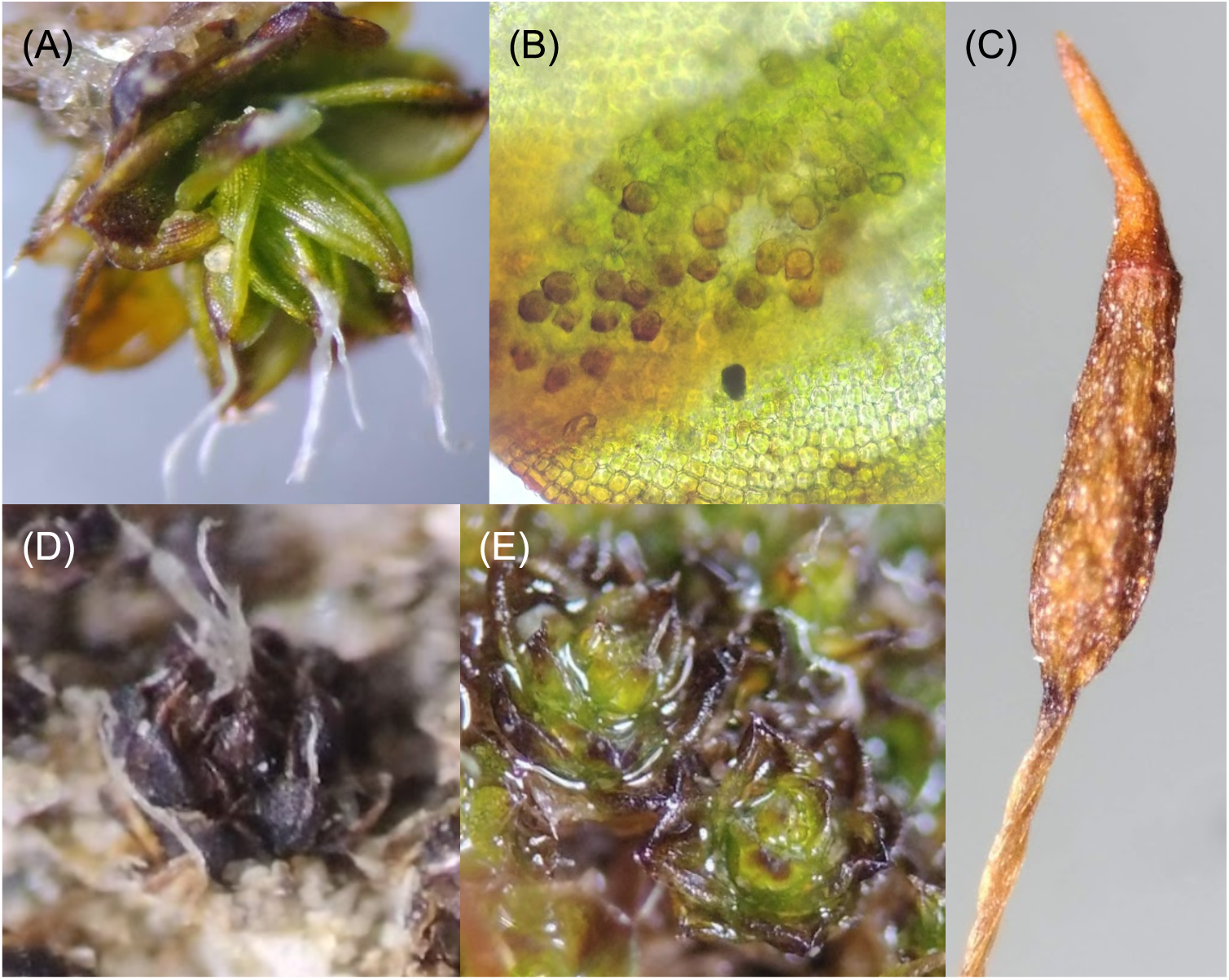
(A) Example *Crossidium squamiferum* gametophyte collected from Anza Borrego Research station. The plant possesses deltoid-ovate leaves with hyaline hairs and adaxial papillae (B) as well as a cylindrical capsule (C) with a long-rostrate operculum. Gametophytes before (C) and after wetting (D) of mixed communities of *Crossidium spp*. collected demonstrate poikilohydry. Green leaves indicate the plant’s health.

**Table 1:** Site coordinates and moss taxonomy.

| Site | Full site name | Climate | Aridity | Moss family | Moss species | Soil type | Longitude | Latitude |
| --- | --- | --- | --- | --- | --- | --- | --- | --- |
| TP | Torrey Pines Site 1 | Coastal | Semi arid | Pottiaceae | *Trichostomopsis australasiae* | Sandstone | -117.2482757 | 32.9421506 |
| TP2 | Torrey Pines Site 2 | Coastal | Semi arid | Pottiaceae | *Vinealobryum brachyphyllum* | Sandy | -117.2482757 | 32.9421506 |
| AB | Anza Borrego Research Center | Colorado Desert | Arid | Pottiaceae | *Crossidium spp. (squamiferum dominant)* | Sandy | -116.3895333 | 33.2407889 |
| ODLO | Oasis De Los Osos Preserve | Colorado Desert | Arid | Pottiaceae | *Trichostomopsis australasiae* | Granitic | -116.68908 | 33.89182 |
| GMT | Granite Mountains Research Station | Mojave Desert | Hyper arid | Grimiaceae | *Grimia sp.* | Granitic | -115.6297361 | 34.7804861 |
| CIMA | Cima Volcanic Field | Mojave Desert | Hyper arid | Pottiaceae | *Trichostomopsis australasiae* | Volcanic | -115.8699722 | 35.1997917 |

### Metabarcode sequencing

Moss biocrust samples were subdivided into three fractions: dirt attached to the biocrust, untreated biocrust, and surface sterilized biocrust. Each field replicate was surface sterilized following the approach of U’Ren et al., 2010. Moss biocrusts were thoroughly rinsed with high pressure water in a soil sieve until as much dirt as possible was removed. Then, they were treated as follows: shaken in 95% ethanol for 30 s, 10% bleach (0.5% NaOCl) for 2 min, and 70% ethanol for 2 min. Finally, they were allowed to dry in a sterile hood at room temperature and stored at −20°C. DNA was extracted using the QIAGEN DNeasy PowerSoil kit (Qiagen, Germantown, MD, USA) from .25 g of either bare soil or moss biocrust associated samples. The manufacturer’s protocol was used with a ten minute extension of bead beating in order to better access DNA within the plant tissue. PCR was performed using ITS1F and ITS2 primers targeting the ITS1 region of fungal ribosomal DNA in accordance with the Earth Microbiome Project (EMP). This will allow future comparisons with the EMP dataset and further characterization of the moss biocrust mycobiome (Thompson et al. 2017). PCR mixtures were as follows: 13 uL PCR grade water (Sigma-Aldrich, St Louis, MO, USA), 10 uL 2X Platinum Hot Start PCR Master Mix (Thermo Fisher Scientific Inc., Waltham, MA, USA), .5 uL of 10uM forward and reverse primer, and 1 uL template DNA for a 25 uL reaction. A C1000 thermal cycler (BioRad, Hercules, CA, USA) was used with the following protocol: 94°C, 1 minute; 94°C, 30 seconds, X35; 52°C, 30 seconds, X35; 68°C, 30 seconds, X35; 68°C, 10 minutes. PCR products were normalized and cleaned using the Invitrogen SequalPrep kit (Invitrogen, Waltham, MA, USA). The resulting pools were quantified using an Agilent 2100 Bioanalyzer and Fragment Analyzer (Agilent Technologies, Santa Clara, CA, USA) and qPCR. The following primer constructs were used for sequencing: Read 1 primer, Read 2 primer, and index primer per the EMP protocol for ITS amplicon sequencing (P. Smith et al. 2018).

Libraries were sequenced using either the illumina MiSeq or NextSeq2000 platform (Illumina, Inc, San Diego, CA, USA) generating 2×300 reads at the Institute for Integrative Genome Biology, Core Facilities, University of California Riverside (http://iigb.ucr.edu). A subset of samples were sequenced to 2×250 read lengths, although bioinformatic processing trimmed all read lengths to 250 to account for this difference. Paired-end sequence reads were deposited in the Sequence Read Archive database associated with BioProject accession number PRJNA1148194.

### Bioinformatics and data analysis

Demultiplexed samples were processed and analyzed using AMPtk version 1.60 (Palmer et al. 2018) (https://github.com/nextgenusfs/amptk). Paired-end reads were merged using VSEARCH (version 2.28.1) (Rognes et al. 2016). Primer sequences were removed with a mismatch of two base pairs allowed and sequences were trimmed to a maximum of 250 base pairs, with reads less than 100 bp discarded. 6,972,409 valid output reads remained after preprocessing.

Sequences were quality filtered to an expected error parameter of 1.0 resulting in a remainder of 6,056,727 reads. The DADA2 pipeline was used to cluster the reads passing these Quality Control filters into Amplicon Sequence Variants (ASVs) (Callahan et al. 2016). The use of ASVs as opposed to OTUs allows for higher resolution comparisons in community composition between samples and will facilitate future comparisons with other datasets. After a total of 1051 *de novo* and reference chimeras were removed using VSEARCH (version 2.28.1) and sequence orientations were validated, a total of 10,413 ASVs remained (Rognes et al. 2016). These ASVs were assigned taxonomy with AMPtk using the UNITE v9.3 database (Abarenkov et al. 2024). The AMPtk default hybrid taxonomic assignment approach was used, which calculates a consensus last common ancestor based on the results of a Global Alignment hybrid taxonomy assignment. This method retains the most specific taxonomic string when results between the two methods do not conflict (Palmer et al. 2018). Finally, a multiple sequence alignment was performed using MAFFT (v7) across all ASVs and a tree was constructed from the alignment using FastTree (v2.1) (Katohand Standley 2013, Price et al. 2010). Clade labels were assigned according to the name of type species occurring, relying on NCBI taxonomy and MaarjAM databases.

### Statistical data analysis and visualization

Data analyses were performed using R version 4.4 (R Core Team 2024), Rstudio version 1.4.1103-4 (RStudio Team 2023), and Phyloseq (McMurdie and Holmes 2013). All samples were rarefied to 7,800 reads. This resulted in the drop of a single sample. Alpha diversity was compared using an ANOVA test using the ‘anova’ function, and a Tukey test with the ‘TukeyHSD’ function in base R. Beta Diversity was calculated using Bray-Curtis dissimilarity as well as unweighted UniFrac phylogenetic distance and compared with a PERMANOVA test using the ‘adonis’ function of the vegan package (v 2.6-7) of R (Oksanen et. al 2001). Venn diagrams were created using the MiscMetabar function ‘ggvenn_pq’ (Taudière, 2023). Phylogenies with heatmaps were visualized using the ggtree function ‘Gheatmap’ and relative abundance was log normalized using base R (Xu et al. 2022). Clades were labeled based on clustering with NCBI type species and the MaarjAM database (Öpik et. al, 2010). All analysis pipelines can be found in the following github repository: https://github.com/stajichlab/moss_mycobiome.

### Microscopy

AMF colonization was visualized using the methodology of Hanke and Rensing, 2010. Briefly, half a milliliter of moss tissue stored at −20°C from each site was rinsed thoroughly with high pressure water in a fine strainer to remove all dirt. Samples were bleached in 10% KOH at 95°C for ten minutes. Then, tissue was acidified in 5% HCl at room temperature for 3 minutes. Samples were then stained using 1% trypan blue for ten minutes at 95°C. Samples were de-stained using 1% chloral hydrate for 30 minutes at room temperature. This step was repeated overnight. Samples were mounted on a slide in glycerol and viewed using an OMAX M83EZ Series Trinocular Compound Microscope (AMscope, Irvine, CA, USA). Unfortunately, the lack of remaining material after DNA extraction meant that only three sites could be surveyed: TP, TP2, and ODLO (table 2).

**Table 2:** Glomeromycotina structures visualized in moss tissue across sites.

| Site | Moss species | Coenocytic hyphae | Hyphal coils | Vesicles | Hyphopodia | Intracellular colonization | Intracellular branching |
| --- | --- | --- | --- | --- | --- | --- | --- |
| ODLO | *Trichostomopsis australasiae* | Stems and leaves | Stems and leaves | None | Stems and leaves | Stems and leaves | None |
| TP | *Trichostomopsis australasiae* | Stems and leaves | Leaves | Leaves | Stems and leaves | Stems and leaves | Leaves |
| TP2 | *Vinealobryum brachyphyllum* | None | None | None | None | None | None |

## Results

### Moss associated Glomeromycotina are distinct from the adjacent bare soil

Two main clades of Glomeromycotina were found to be associated with moss biocrusts: *Rhizophagus* and *Glomus* (Figure 3). In contrast, *Funneliformis* and *Septoglomus* were primarily found in samples from bare soil. A clear distinction in the phylogenetic distribution of AMF recovered in bare soil and moss crust samples was apparent, especially after surface sterilization of the moss samples. A semi-consistent set of Glomeromycotina ASVs were recovered from bare soil across sites and these were distinct from the Glomeromycotina found in the moss biocrusts (Figure 3). In total, 20 Glomeromycotina ASVs were shared between bare soil and the moss crust while 74 were found only in association with mosses (Figure 5C). When comparing moss crust samples to bare soil samples, there was a statistically significant difference in Bray-Curtis beta diversity (PERMANOVA p = .009, R2 = 0.05459). However, there was no difference in observed alpha diversity between moss crust samples and bare soil (Figure S1).

**Figure 3:**
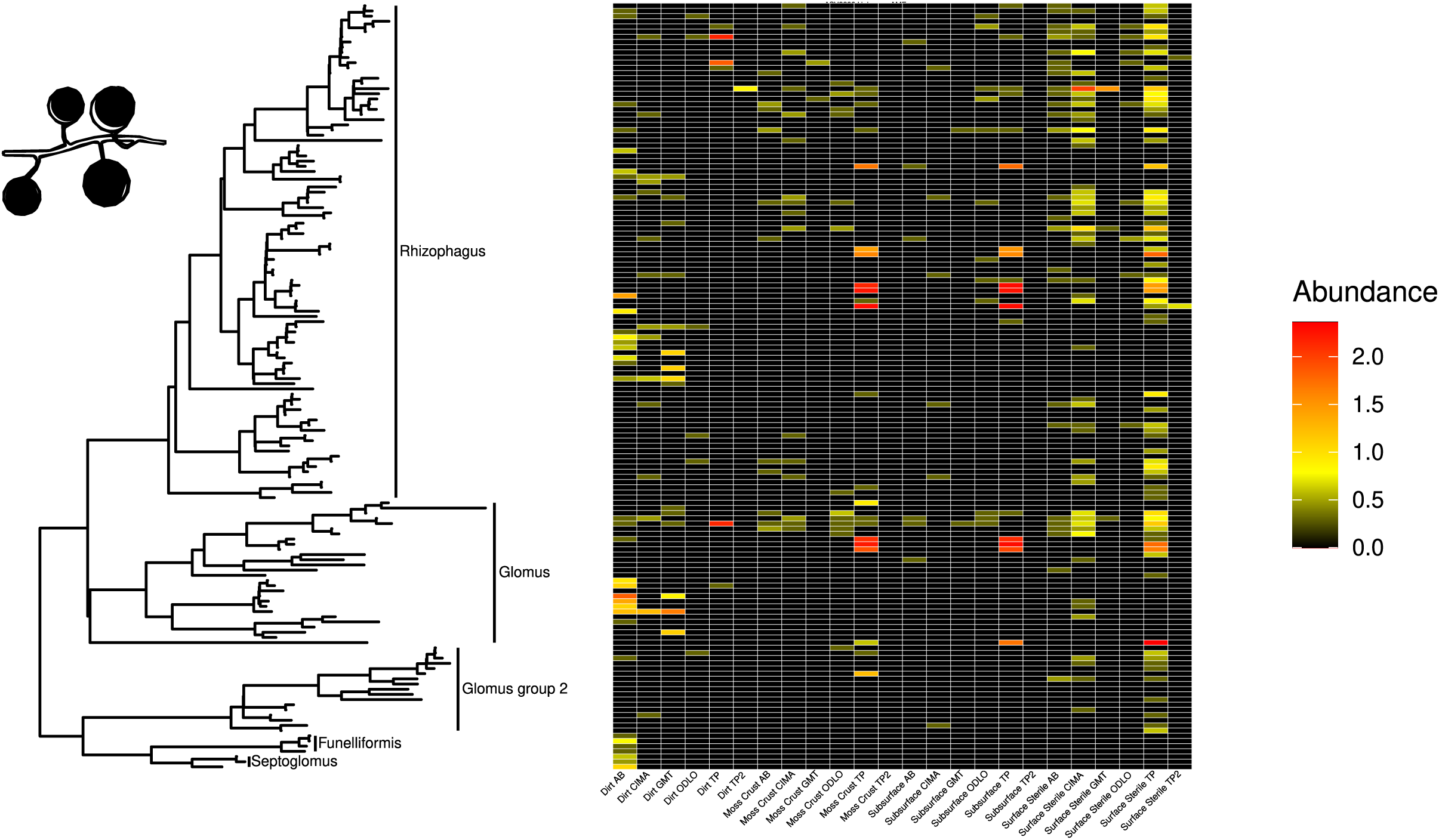
Normalized Glomeromycotina relative abundance by site and substrate. Black, yellow, orange and red represent low, medium, medium high, and high normalized abundance respectively. Surface sterilization results in a similar set of moss-associated Glomeromycotina in TP, CIMA, AB, and ODLO. Bare soil (dirt) samples possess a phylogenetically distinct palette Glomeromycotina ASVs compared to all moss crust samples. Clades were named based on clustering with NCBI type species and the MaarjAM database. TP = Torrey Pines, ODLO = Oasis De Los Osos Reserve, AB = Anza Borrego Research Station, CIMA = CIMA Volcanic Field, GMT = Granite Mountains Research Center. Subsurface = dirt connected to the biological soil crust. All samples were rarefied to 7800 reads.

### Surface sterilization reveals diverse Glomeromycotina

Alpha diversity of AMF was not enriched after surface sterilization (Figure S1), and Bray-Curtis beta diversity was unchanged by sterilization (PERMANOVA p = 0.683, R2 = 0.03056). However, surface sterilization of moss biocrusts increased the relative abundance of a diverse set of Glomeromycotina (Figure 3). A total of 28 Glomeromycotina ASVs were found exclusively after the surface sterilization treatment (Figure 5D). 23 ASVs were shared between sterilized and untreated moss biocrusts, indicating taxa that are not dependent on surface sterilization for recovery by metabarcoding. 5 unique Glomeromycotina ASVs were found in the unsterilized moss crusts.

### Fungi morphologically similar to Glomeromycotina form intracellular branches in moss leaves

Key signs of mycorrhizal fungi colonizing moss were observed at TP in *T. australasiae* (Figure 4); these include vesicles (4C), hyphal coils (4A), coenocytic hyphae (4D), hyphopodia (4E), and intracellular branching (4A). These results are corroborated by the observation that TP *T. australasiae* had the highest relative abundance of Glomeromycotina by an order of magnitude (ANOVA, p = 0.05) (Figure 5B). ODLO *T. australasiae* also contained coenocytic hyphae, intracellular colonization, and hyphal coils, but no vesicles or intracellular branching was observed (Table 2). These results suggest that climate may control the formation of intracellular branching and vesicles within moss tissue. No fungal hyphae were observed after staining TP2 *Vinealobryum brachyphyllum* (Table 2), which is consistent with culture independent survey results.

**Figure 4:**
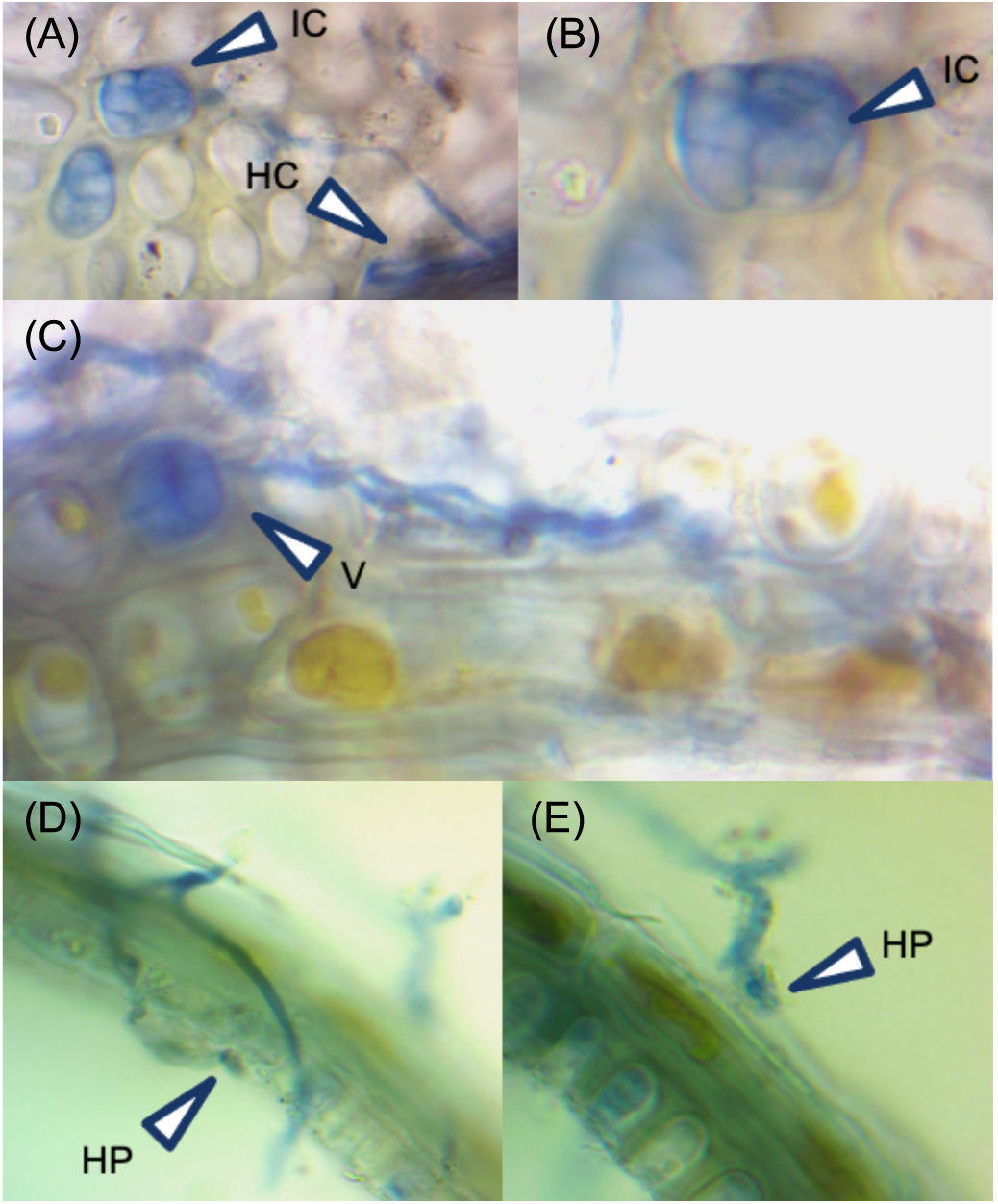
Putative Glomeromycotina colonization of moss at TP observed with trypan blue stain. (A) Leaf cells demonstrating coenocytic hyphae with hyphal coiling and extensive intracellular branching (60X) (B) (100X) view of the colonized cell from panel A. (C) Vesicles posessing coenocytic hyphae observed towards the center of the leaf (100X). (D, E) Hyphopodia observed at the leaf stem (60X). IC = Intracellular colonization, HC = Hyphal coil, HP = Hyphopodium, V = Vesicle.

**Figure 5:**
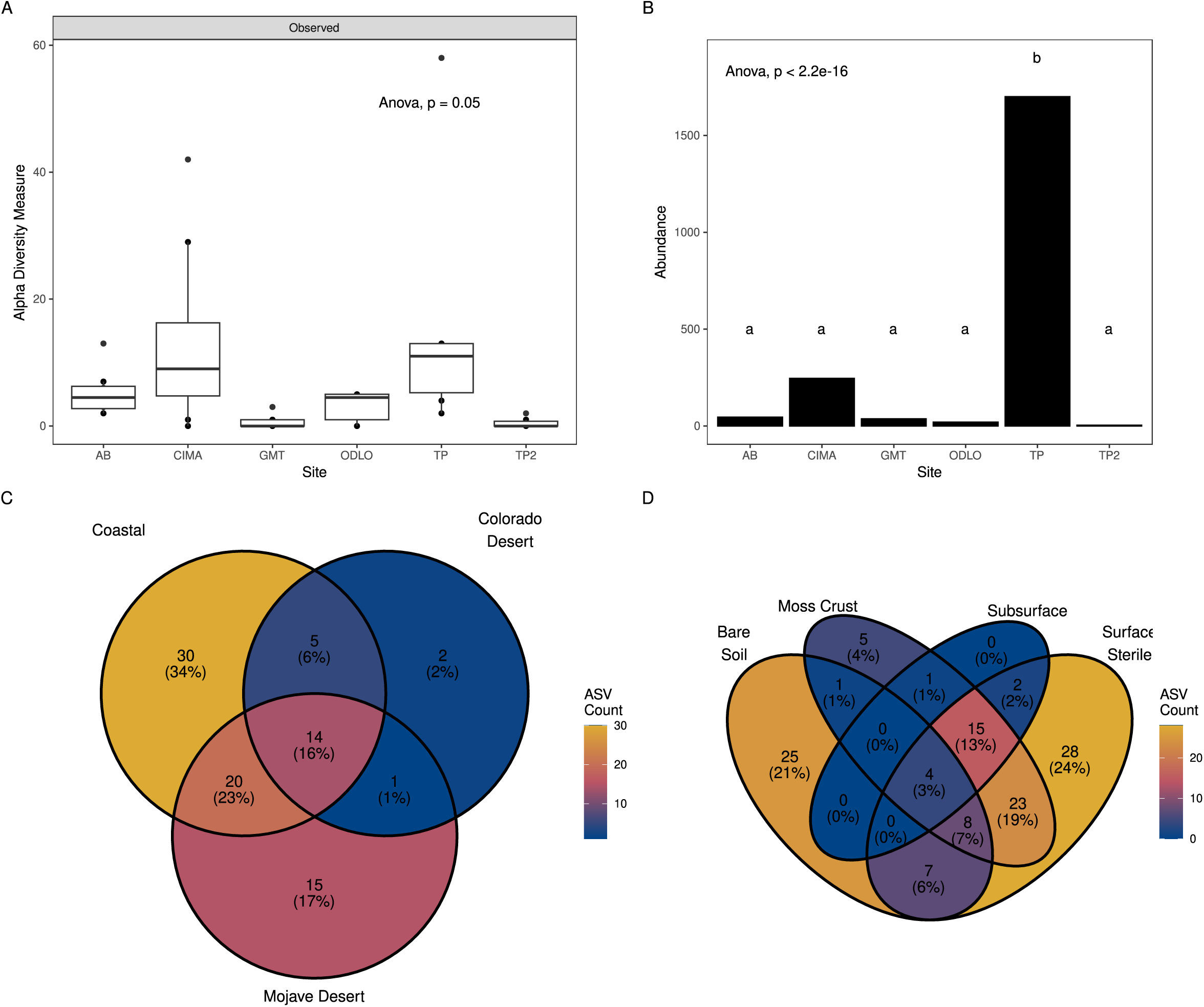
(A) Observed alpha diversity of Glomeromycotina by site within both surface sterilized and untreated moss crust. (B) Relative abundance of Glomeromycotina by site in surface sterilized and untreated moss crust, presented as summed read counts per Glomeromycotina ASV. Letters indicate Tukey HSD significance (p < 0.05). (C) Venn diagram of Glomeromycotina ASV counts by climate post-surface sterilization. (D) Venn diagram illustrating Glomeromycotina ASV counts by substrate. “Subsurface” denotes dirt connected to the biological soil crust TP = Torrey Pines, ODLO = Oasis De Los Osos Reserve, AB = Anza Borrego Research Station, CIMA = CIMA Volcanic Field, GMT = Granite Mountains Research Center. All samples were rarefied to 7800 reads.

### Glomeromycotina-moss associations differed between climates

A comparison of Bray-Curtis beta diversity of Glomeromycotina across sites revealed significant differences in community composition within moss crust and sterile moss crust samples (PERMANOVA, p = 0.003, R² = 0.26897) (Figure S3). A similar trend was apparent for observed alpha diversity (ANOVA, p < 0.05) (Figure 5A). Notably, despite TP and CIMA sites sharing the same host species (*T. australasiae*), their beta diversity patterns were distinct in moss crust and sterile moss crust samples (PERMANOVA, p < 0.05, R² = 0.18075) (Figure 6A). TP exhibited greater UniFrac phylogenetic dissimilarity of arbuscular mycorrhizal fungi (AMF) between samples compared to CIMA. A similar pattern was observed across all collected moss crust samples: semi-arid climates displayed higher UniFrac phylogenetic dissimilarity between samples than arid and hyper-arid climates, though different moss species were present at some sites and this difference was not statistically significant (PERMANOVA, p = 0.088, R² = 0.1307) (Figure 6B). Surface-sterilized samples revealed a core set of 14 moss-associated Glomeromycotina ASVs present across all sites, with 20 ASVs common to both the Mojave Desert and coastal habitats (Figure 5C). Coastal samples exhibited the highest number of unique ASVs (30), followed by the Mojave Desert (15), while the Colorado Desert had the fewest unique endophytic AMF ASVs (2) (Figure 5C).

**Figure 6:**
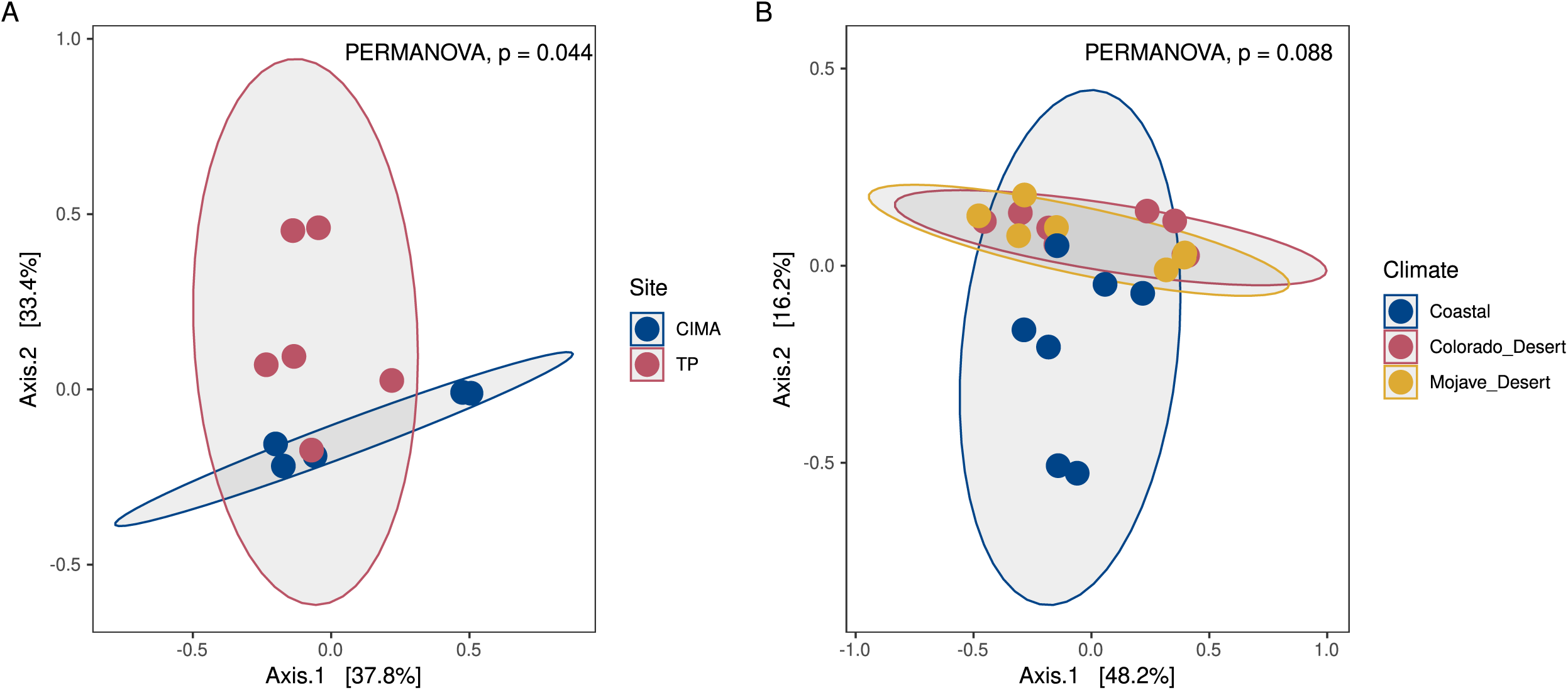
(A) Unweighted unifrac phylogenetic beta diversity of Glomeromycotina between CIMA and TP within both surface sterilized and untreated moss crust. Note: CIMA and TP contain the same moss species, *Trichostomopsis australasiae* (B) Beta diversity of Glomeromycotina climate, across all moss species including surface sterilized and untreated moss crust. TP = Torrey Pines, CIMA = CIMA Volcanic Field. All samples were rarefied to 7800 reads.

## Discussion

Our survey results illustrate several key findings: (i) Surface sterilization unveils diverse AMF; (ii) moss-associated AMF diversity varies by local climate and site; (iii) Moss-associated AMF differ from adjacent bare soil; (iv) Cells morphologically similar to Glomeromycotina can form intracellular branches in *T. australasiae* leaves.

### Surface sterilization reveals diverse Glomeromycotina

Only a handful of studies have characterized the diversity of Glomeromycotina within mosses (Zhang et al. 2021, Zhang and Guo 2007, Valdés et al. 2023). However, to our knowledge, only one study has utilized surface sterilization to characterize the moss endophytome in a culture independent survey (Zhang et al. 2021). This methodological gap may explain the limited recovery of Glomeromycotina from moss samples in previous studies (U’Ren et al., 2010, Zhang et al. 2021, Pombubpa et al. 2020). Another explanation for low Glomeromycotina recovery in many moss biocrust surveys may be the use of primers not suited to early divergent fungi. Future studies should couple the use of AMF specific primers and surface sterilization to further characterize the moss biocrust mycobiome.

### Moss biocrust associated Glomeromycotina diversity is associated with local climate

Sampling the moss species *T. australasiae* across multiple sites allowed us to explore patterns of host specificity and biogeographic diversity. At TP, *T. australasiae* exhibited greater UniFrac phylogenetic dissimilarity of AMF between samples compared to CIMA, and TP also showed an order of magnitude higher relative abundance of AMF. This suggests that semi-arid climates, such as TP, may be more suitable to diverse moss-associated Glomeromycotina communities than hyper-arid sites like CIMA. However, CIMA still exhibited relatively high alpha diversity, indicating the presence of niche-specialized, low-abundance Glomeromycotina in this hyper-arid environment. This is noteworthy since Glomeromycotina are generally not associated with hyper-arid climates.

The observed differences in the biocrust mycobiome across the aridity gradient point to potential shifts in Glomeromycotina composition and relative abundance as climate change progresses. These shifts may contribute to the high vulnerability of moss biocrusts to climate change, as shown in previous studies (Ferrenberg et al., 2015). It is known that AMF symbiosis in desert plants is influenced by aridity, which can impact plant growth outcomes (Chávez-González et al., 2024). Overall, these results suggest that local climate may play a role in shaping the diversity and abundance of moss-associated Glomeromycotina, with important implications for understanding the impact of climate change on biocrusts. It is important to note that this study did not take into account variations in soil pH, nutrient content, salinity, UV light intensity, seasonality, and plant density which may impact the composition of the moss biocrust mycobiome. Future studies should take these variables into account.

### Moss associated Glomeromycotina are distinct from the adjacent bare soil

Previous investigations into moss-associated arbuscular mycorrhizae have not made a clear differentiation between the fungal communities within the moss tissues (endophytome) and those present in the surrounding soil (Zhang et al., 2021; Zhang and Guo, 2007; Valdés et al., 2023). This distinction is crucial, given previous hypotheses suggesting that Glomeromycotina function as saprotrophs on mosses, possibly derived from neighboring host plants (Zhang and Guo, 2007). However, such claims lack empirical support. The previously established specificity of fungal (non-AMF) associations with different moss species implies a non-random nature of these interactions (Zhang et al., 2021). In this study, we observed phylogenetic disparities between the AMF communities inhabiting moss biocrusts and those present in the adjacent bare soil, underscoring the specificity of bryophyte-fungal relationships. Surface sterilization procedures unveiled a distinct palette of Glomeromycotina, further emphasizing the phylogenetic uniqueness of arbuscular mycorrhizal fungi (AMF) within mosses compared to the surrounding soil.

### Glomeromycotina can form intracellular branches in Trichostomopsis australasiae leaves

Multiple studies have demonstrated the intracellular colonization of mosses by Glomeromycotina (Zhang and Guo, 2007; Hanke and Rensing, 2010; Valdés et al., 2023). However, none of these investigations have provided evidence of intracellular *branching* in healthy cells, which may indicate nutrient exchange. To our knowledge, our findings present the first documented images of intracellular branching within healthy moss cells. This challenges the traditional understanding that mosses are entirely asymbiotic, as intracellular branching within healthy plant cells is a characteristic feature of symbiotic interactions (Field et al., 2015; Hoysted et al., 2023). Future research efforts should determine whether Glomeromycotina fungi establish intracellular branches in *T. australasiae* to facilitate nutrient exchange. Comparative analyses of *T. australasiae*-Glomeromycotina associations and mycorrhizal symbioses with more evolutionarily advanced plants, such as angiosperms, may provide valuable insights into the evolutionary trajectory of plant-fungal symbiotic relationships and offer insights into the symbioses that enabled the conquest of land by plants.

## Conclusions

Our study revealed several key findings. First, we identified a widespread community of Glomeromycotina in healthy moss crusts across diverse regions, including the Mojave Desert, Colorado Desert, and California Coast. Our survey also revealed novel associations between *Rhizophagus* and *Glomus* species with mosses including *T. australasiae*. Phylogenetic analysis highlighted distinct evolutionary relationships between Glomeromycotina associated with mosses and those found in bare soil, underscoring the specificity of these moss-fungal associations. Moreover, staining of moss tissue revealed intracellular branching of Glomeromycotina within healthy moss cells, suggesting symbiotic interactions beyond saprotrophy.

We also observed shifts in the composition of moss-associated Glomeromycotina along the aridity gradient, particularly in the host species *T. australasiae*. Structures such as vesicles and intracellular branches were exclusively observed in semi-arid *T. australasiae*, with no such structures found in *T. australasiae* from arid climates. These findings indicate that climate may influence the Glomeromycotina mycobiome in biocrusts, and suggest that these communities may shift in composition and abundance under climate change. This has significant implications for understanding how climate change could affect the health and functionality of biocrusts. Future studies should further explore whether the associations between *T. australasiae* and *Glomus* or *Rhizophagus* species represent true symbioses. Our findings provide valuable insights into ancient plant-fungal interactions and lay the groundwork for future studies exploring the mycobiomes of the approximately 10,000 known moss species. They also suggest fungal taxa which may serve as inoculants to help promote biocrust resilience to climate change.

## Supporting information

Includes all supplemental figures

## Acknowledgements

We extend our gratitude to the Mojave National Preserve (Permit MOJA-2023-SCI-0043), as well as the University of California Natural Reserves Anza Borrego Desert Research Center (Application #52877) and Oasis De Los Osos (Application #52886). We also thank Torrey Pines State Natural Reserve (permit application number 23-630-03) for granting us permission to conduct this research. Our thanks go to Cassie Ettinger, Xinzhan Liu, Mark Yacoub, Cheng-Hung Tsai, and especially Michael Remke for their helpful feedback and suggestions. We thank Jessica Wu-Woods, Leila Shadmani, and Sadikshya Sharma for their assistance in laboratory logistics and management. Finally, we acknowledge the use of the following Creative Commons licensed photos: “Vista of Anza Borrego” by John Fowler, used under CC BY-SA 3.0 (cropped from original); “View from Guy Fleming Trail” by Remember to Breathe, used under CC BY-NC-ND 2.0 (cropped from original); and “Granite Mountains” by Unknown, used under Public Domain Mark 1.0 Universal. Analyses were performed on the UC Riverside High Performance Computing Cluster supported by grants from the National Science Foundation (DBI-1429826 & DBI-2215705) and the National Institutes of Health (NIH) (S10-OD016290). JES is a CIFAR fellow in the program Fungal Kingdom: Threats and Opportunities and was supported by USDA (National Institute of Food and Agriculture Hatch project CA-R-PPA-211-5062-H).

## Author Contributions

JES and KHK conceived of and planned the study. Field sampling was conducted by KHK. Mosses were identified by C. Coshland. DNA extraction, PCR, normalization, and library preparation were conducted by KHK. Bioinformatics analyses were conducted by KHK and JES. Statistical data analysis and visualization was conducted by KHK. Project administration and funding acquisition were carried out by JES. Manuscript was written by KHK with contributions from JES and C. Coleine.

