## Supplementary material for "Novel Glomeromycotina-Moss Associations Identified in California Dryland Biocrusts": Includes all supplemental figures

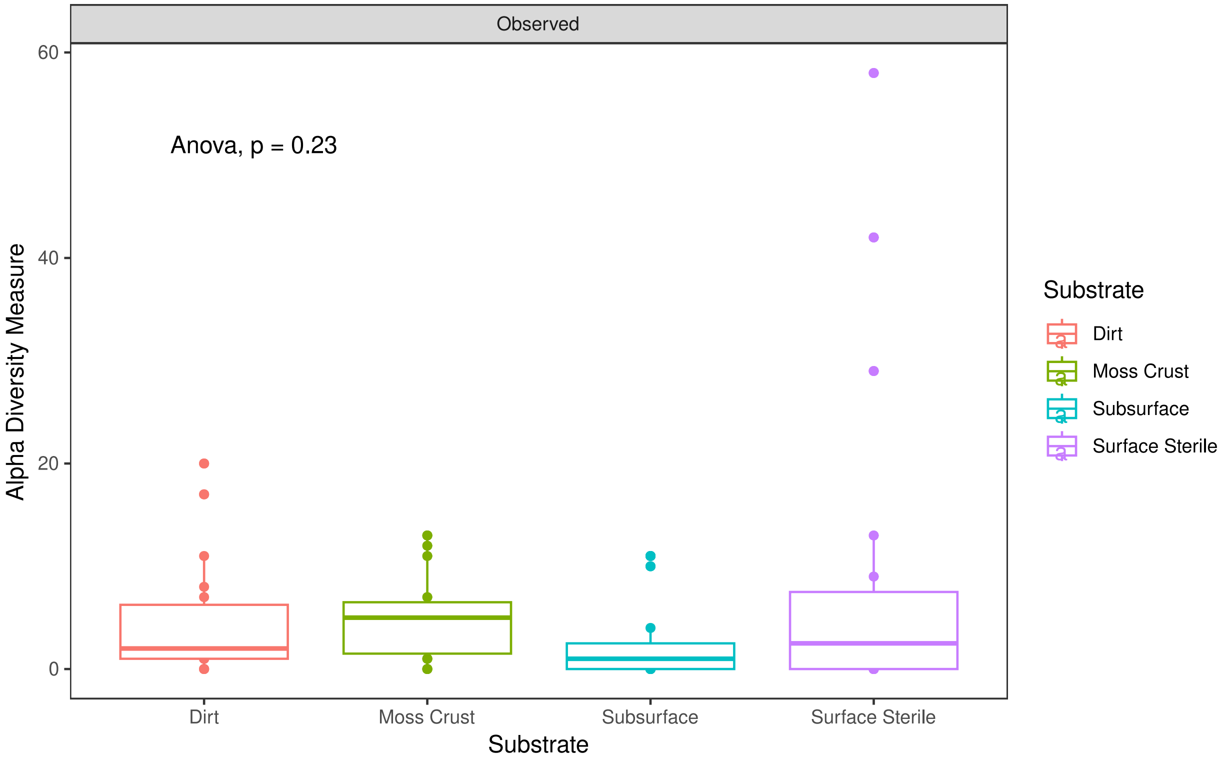


**Figure S1:** Glomeromycotina alpha diversity stratified by substrate.

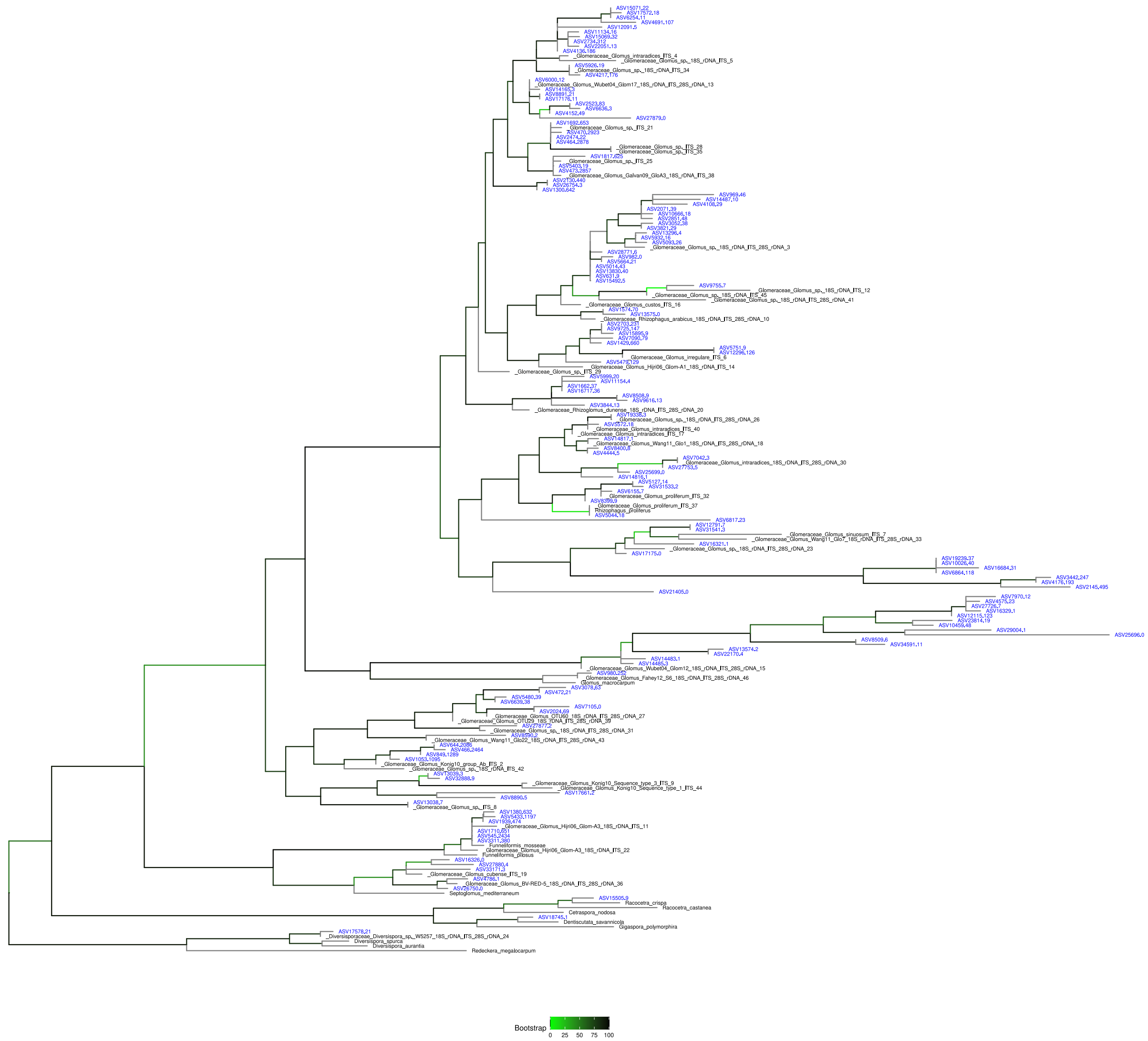


**Figure S2:** Phylogenetic tree of Glomeromycotina ASVs (blue), NCBI type species and closest hits in the MaarjAM database (black). Bootstrap values are represented as a branch color gradient from low (green) to high (black).

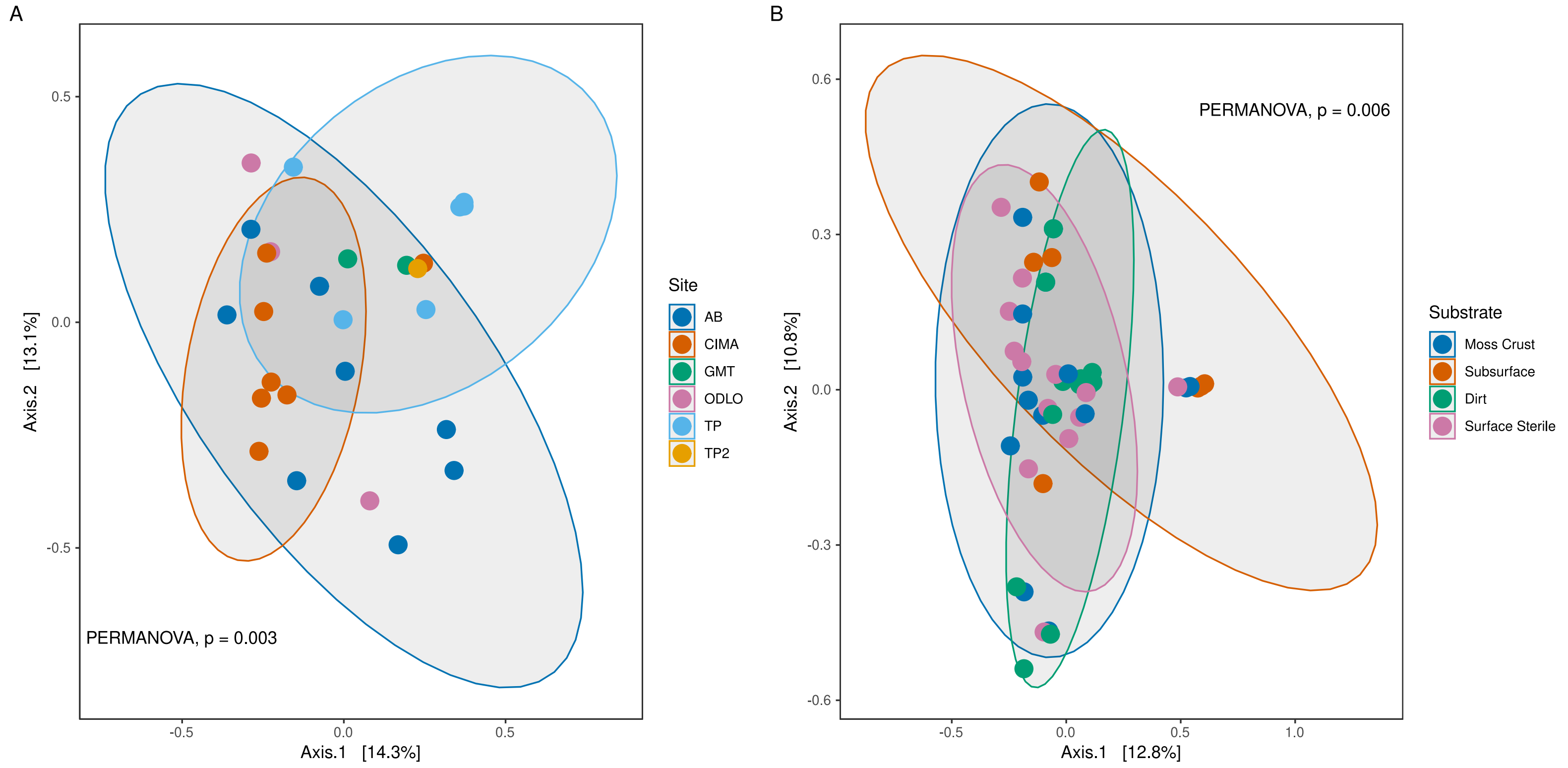


**Figure S3:** (A) Bray-Curtis beta diversity of Glomeromycotina by site for surface sterilized and untreated moss crust. (B) Bray-curtis beta diversity of Glomeromycotina by substrate. TP = Torrey Pines, ODLO = Oasis De Los Osos Reserve, AB = Anza Borrego Research Station, CIMA = CIMA Volcanic Field, GMT = Granite Mountains Research Center. All samples were rarefied to 7800 reads.
